# The Histidine Kinase NahK Regulates Denitrification and Nitric Oxide Accumulation through RsmA in *Pseudomonas aeruginosa*

**DOI:** 10.1101/2024.06.07.597968

**Authors:** Danielle Guercio, Elizabeth Boon

**Affiliations:** Graduate Program in Molecular and Cellular Biology, Stony Brook, NY; Department of Chemistry, Stony Brook, NY; Institute of Chemical Biology and Drug Discovery; Stony Brook University, Stony Brook, NY

**Author notes:** Corresponding author: Elizabeth M. Boon.

**Keywords:** nitric oxide, biofilm, NosP, NahK, RsmA, denitrification, quorum sensing, cell elongation

## Abstract

*Pseudomonas aeruginosa* have a versatile metabolism; they can adapt to many stressors, including limited oxygen and nutrient availability. This versatility is especially important within a biofilm where multiple microenvironments are present. As a facultative anaerobe, *P. aeruginosa* can survive under anaerobic conditions utilizing denitrification. This process produces nitric oxide (NO) which has been shown to result in cell elongation. However, the molecular mechanism underlying this phenotype is poorly understood. Our laboratory has previously shown that NosP is a NO-sensitive hemoprotein that works with the histidine kinase NahK to regulate biofilm in *P. aeruginosa*. In this study, we identify NahK as a novel regulator of denitrification under anaerobic conditions. Under anaerobic conditions, deletion of *nahK* leads to a reduction of growth coupled with reduced transcriptional expression and activity of the denitrification reductases. Further, during stationary phase under anaerobic conditions*, ΔnahK* does not exhibit cell elongation, which is characteristic of *P. aeruginosa*. We determine the loss of cell elongation is due to changes in NO accumulation in *ΔnahK*. We further provide evidence that NahK may regulate denitrification through modification of RsmA activity.

**Importance:** *P. aeruginosa* is an opportunistic multi-drug resistance pathogen that is associated with hospital acquired infections. *P. aeruginosa* is highly virulent, in part due to its versatile metabolism and ability to form biofilms. Therefore, better understanding of the molecular mechanisms that regulate these processes should lead to new therapeutics to treat *P. aeruginosa* infections. The histidine kinase NahK has been previously shown to be involved in both NO signaling and quorum sensing through RsmA. The data presented here demonstrate that NahK is responsive to NO produced during denitrification to regulate cell morphology. Understanding NahK’s role in metabolism under anaerobic conditions has larger implications in determining Nahk’s role in a heterogeneous metabolic environment such as a biofilm.

## Introduction

The opportunistic pathogen *P. aeruginosa* is a major contributor to ventilator-associated pneumonia, catheter-associated urinary tract infections, and burn wound infections (1-3). In particular, cystic fibrosis (CF) patients are highly susceptible to *P. aeruginosa* lung infections. *P. aeruginosa* thrive within the thick microaerobic mucus layers in the lungs of CF patients, resulting in chronic infections and damage to epithelial tissue (4). *P. aeruginosa* are facultative aerobes; under anaerobic conditions, they utilize nitrate or nitrite as alternative electron acceptors in a process termed denitrification (5). During denitrification, nitrate is reduced to nitrogen through the sequential activity of the reductases nitrate reductase (NAR), nitrite reductase (NIR), NO reductase (NOR), and nitrous oxide reductase (NOS) (6), thus supporting growth in the absence of oxygen.

Under anaerobic conditions, the principal transcriptional regulator for genes associated with denitrification, including the transcription factor DNR, is ANR. DNR then promotes the transcriptional expression of the denitrification reductases and is additionally activated by NO, an intermediate in the denitrification process produced by NIR activity (7). This is hypothesized to be a mechanism to prevent accumulation of toxic levels of NO within the cell during denitrification (8). This is important because at high concentrations, NO disrupts iron-sulfur clusters and other iron-binding proteins, nitrosylates cysteines, and causes lipid peroxidation and DNA strand breaks, leading to changes in cell membrane permeability and membrane potential and cell death (9-10).

In addition to being regulated by NO concentrations through DNR, denitrification reductases are transcriptionally regulated by the *las*, *rhl*, and *pqs* quorum sensing networks (11-12). The *las* and *rhl* systems transcriptionally downregulate the denitrification reductases; strains that lack either of the response regulators *lasR* or *rhlR* have been shown to have an increase in denitrification activity due to increased expression of the reductases (11). In PAO1 *ΔrhlR,* increased expression of NAR and NIR contribute to an overproduction of NO, leading to cell death in anaerobic biofilms (13). Further, the quorum sensing autoinducer PQS has been shown to reduce anaerobic growth by directly inhibiting the activity of the denitrification reductases NAR, NOR, and NOS through active-site iron chelation; in contrast PQS promotes the activity of NIR (12). However, in fully anaerobic conditions, PQS is not produced, because its production relies on the monooxygenase PqsH, suggesting that its role in dentification may only be relevant in microaerobic conditions or the transition between aerobic and anaerobic respiration (11-12,14).

NO is well documented as a signal for biofilm dispersal. In *P. aeruginosa,* the NO sensing hemoprotein NosP has been shown to be necessary for NO-mediated biofilm dispersal by inhibiting a co-cistronic histidine kinase NahK (15-16). NahK is part of the GacS multi-kinase network (MKN) (17). NahK transfers phosphate to the histidine-containing phosphotransfer protein HptB (18). When HptB is unphosphorylated, it plays a role in regulating transcription of the small regulatory RNA *rsmY* (19). *rsmY* inhibits translation of the RNA-binding protein RsmA, a post-transcriptional regulator of 100s of genes, which is thought to be the master switch between biofilm and planktonic growth, promoting bacterial motility and inhibiting quorum sensing and biofilm formation (20).

Recently we reported that a *nahK* deletion strain dramatically over produces the molecule pyocyanin through mis-regulation of the PQS quorum sensing system (21). Pyocyanin is a redox active small molecule secreted by *P. aeruginosa* to act as an electron shuttle supporting aerobic respiration under the microaerobic conditions in a biofilm (22-23). NahK is not alone as a regulator of PQS in the GacS MKN; it has also been reported that *ΔrsmA*, *ΔgacS* and *ΔretS* overproduce pyocyanin and *ΔrsmY* and *ΔrsmZ* underproduce pyocyanin, relative to wild-type (19, 24). The overproduction of pyocyanin and mis-regulation of quorum sensing in *ΔnahK* (21), as well as its role in NO-mediated biofilm regulation (15), led us to hypothesize that NahK may play a regulatory role in denitrification, as both NO and pyocyanin are essential for survival in microaerobic conditions. Here we describe a novel role for NahK in promoting anaerobic biofilm formation and cell elongation through regulation of intracellular NO accumulation.

## Results

### Deletion of nahK leads to changes in anaerobic growth

To investigate if NahK has a role in regulation of respiration under micro- or anaerobic conditions, first we investigated the role of NahK in growth under anaerobic denitrifying (addition of NaNO_3_) conditions in comparison to a wild-type strain (Figure 1A). Under anaerobic conditions, *ΔnahK* initially grows faster in early exponential phase; after 1 hour of anaerobic growth, the CFU value of *ΔnahK* is higher than wild-type (Figure 1B). However, over time the wild-type strain catches up; the CFUs after four hours of growth and during stationary phase are lower in *ΔnahK* in comparison to wild-type (Figure 1C, 1D). These results suggest that *nahK* alters anaerobic growth and may be required for survival under these conditions.

**Figure 1.**
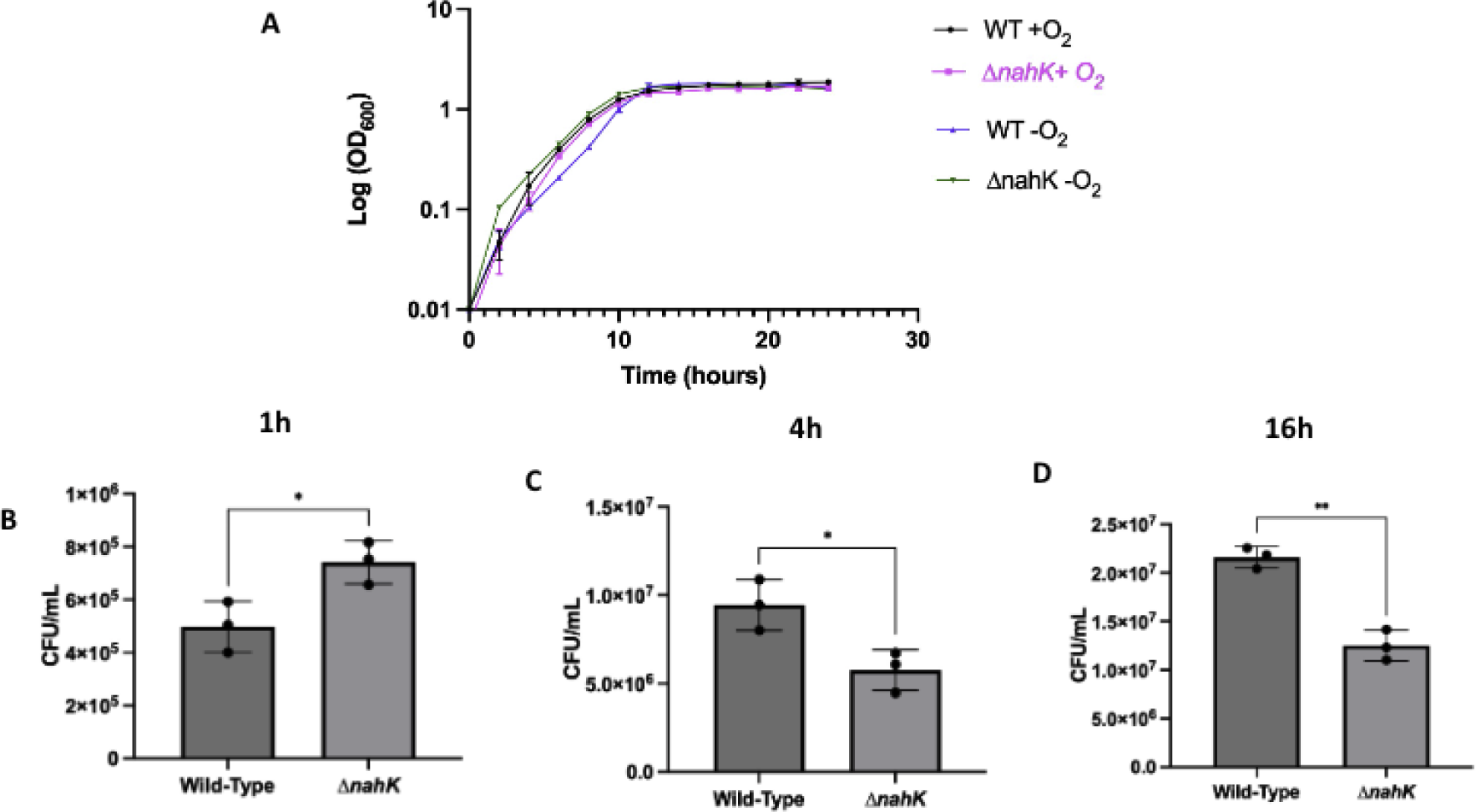
Anaerobic growth is altered in *ΔnahK*. **(A)** Growth curves under aerobic and denitrifying conditions. The average OD readings of 3 independent experiments are plotted as a function of time. **(B)** CFUs after 1 h anaerobic growth. **(C)** CFUs after 4 h of growth under denitrifying conditions. **(D)** CFUs after 16 h of growth under denitrifying conditions. The plotted values are the average ± 1 standard deviation from 3 independent experiments. *p* values were calculated using unpaired, t-tailed t-test comparing ΔnahK OD to Wild-type OD. \**p* ≤ 0.5.

### Denitrification reductases are differentially expressed during exponential and stationary phase

To understand why we observed a growth difference in Δ*nahK*, under anaerobic conditions, we conducted qPCR on the reductase transcripts as well as the transcripts of the denitrification regulators *anr* and *dnr*. During exponential phase (after 4 h), we observed an upregulation in transcription of both *dnr* and the reductases in *ΔnahK*, relative to wild-type, which could explain the increase in initial growth (Figure 2A). Further, during stationary phase (after 16 h), we observed a decrease in expression of the reductases and *dnr* (Figure 2B) in *ΔnahK*, relative to wild-type, which could account for the observed reduced growth after early exponential phase. No change in *anr* transcript levels was observed in either condition.

**Figure 2.**
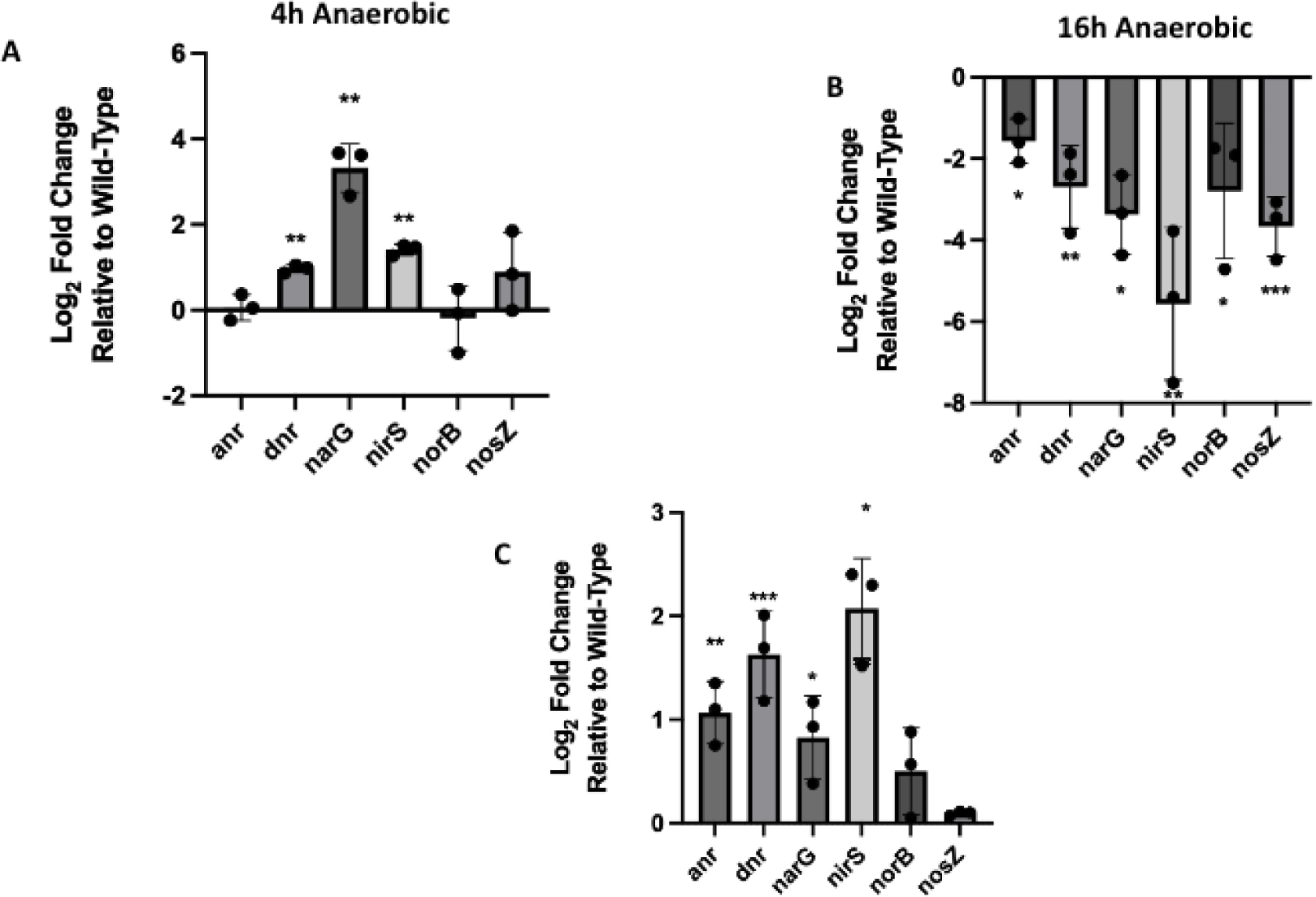
Transcription of the denitrification machinery is mis-regulated in *ΔnahK*. Transcript levels in Δ*nahK*, relative to wild-type, as measured by qPCR for the four denitrification reductases and two transcriptional regulators. Gyrase A (gyrA) was used as a housekeeping gene. Bars above and below the threshold represent up- and down-regulation in Δ*nahK*, relative to wild-type, respectively. The values represent average ± 1 standard deviation of 3 independent experiments. RNA was extracted at several time points and growth conditions: **(A)** after 4 h anaerobic growth (exponential stage); **(B)** after 16 h anaerobic growth (early stationary phase); **(C)** and at an OD of 1.0 during aerobic growth. *p* values were calculated using unpaired, two-tailed t-tests comparing Δ*nahK* ΔCt values to wild-type ΔCt values; n = 3; \**p* value < 0.05, \*\**p* value < 0.01, \*\*\**p* value < 0.005.

Because the increase in reductase expression in Δ*nahK* was observed right after the culture was inoculated, we next wanted to see increased expression of the reductases was occurring under aerobic conditions. Under aerobic conditions, there is an increase in expression of *anr*, *dnr narG*, and *nirS* (Figure 2C) in *ΔnahK*, relative to wild-type. This increased expression could indicate that the anaerobic respiration machinery is mis-regulated in Δ*nahK* and the bacteria are behaving as if they are under microaerobic conditions even during aerobic respiration, and thus at the time of inoculation were primed for anaerobic respiration. This is consistent with inhibition of the *las* and *rhl* quorum sensing systems in *ΔnahK*, as we have previously reported (11).

### Reductase activity is reduced in ΔnahK

Since we observed the greatest differences in expression of the reductases NAR and NIR in Δ*nahK* relative to wild-type, during both exponential and stationary phase, we wanted to determine if we could measure differences in denitrifying enzyme activities under these conditions. To accomplish this, we looked at nitrite production (from nitrate), nitrite utilization, and NO accumulation, each as a function of time, as read outs for the activity of the reductases NAR and NIR.

NAR is responsible for the reduction of nitrate into nitrite, therefore measuring nitrite production was used as a readout for NAR activity. To measure nitrate reduction, we supplemented cultures with nitrate and measured nitrite production (Figure. 3A, 3B). Nitrite is steadily produced during the entire time course in the wild-type strain. However, after 4 h of growth, we observed very little production of nitrite in *ΔnahK* (Figure 3A, 3B).

**Figure 3.**
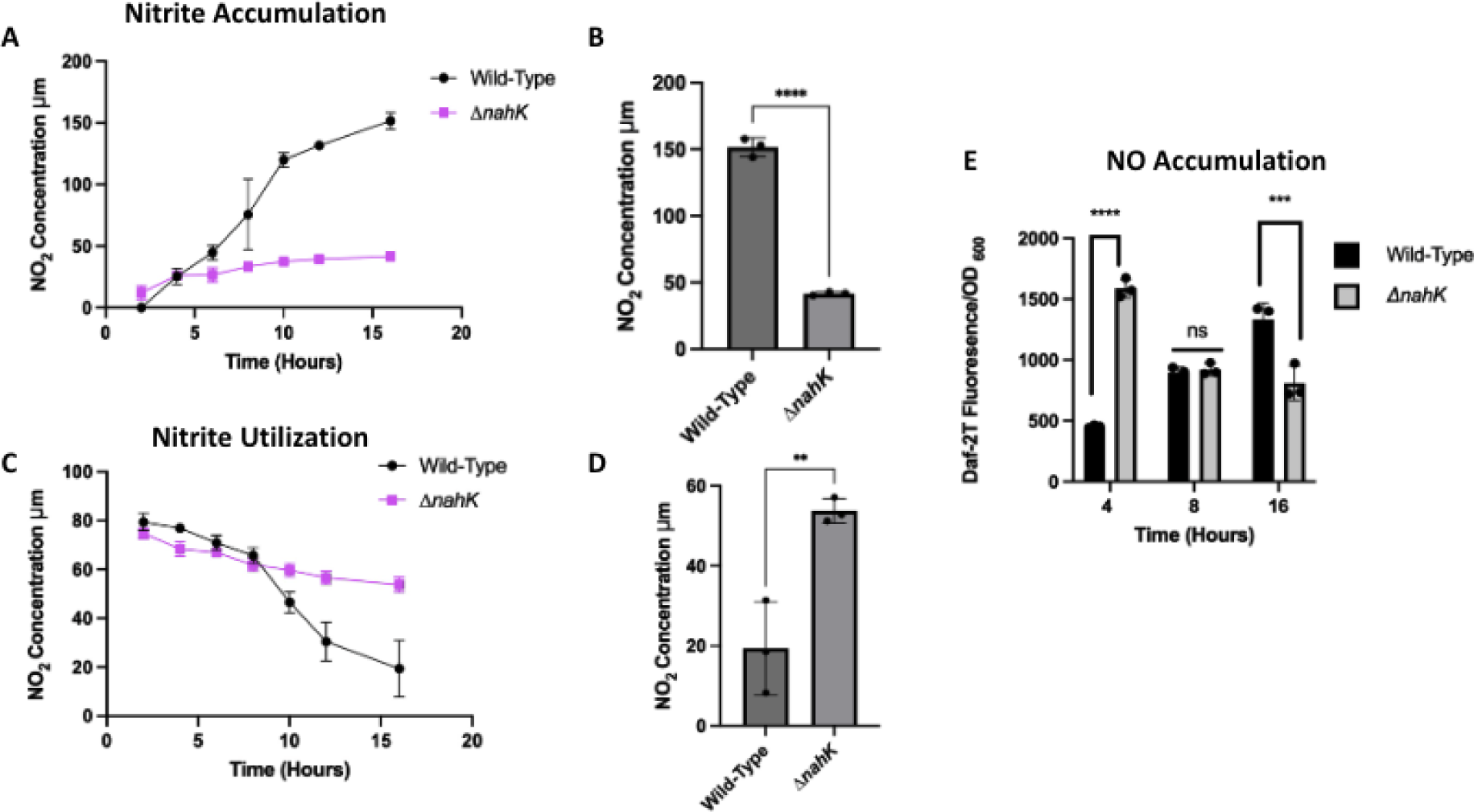
Denitrification activity is mis-regulated in *ΔnahK*. **(A)** Nitrite production by NAR during anaerobic denitrifying conditions, as a function of time. **(B)** Nitrite production quantified after 16 h of anaerobic growth (stationary phase). Initially, *ΔnahK* produces a little more nitrite than wild-type, but over time nitrite production stagnates in the mutant while steadily increasing in the wild-type. **(C)** Nitrite utilization by NIR during anaerobic denitrifying conditions, as a function of time. **(D)** Nitrite utilization quantified after 16 h of anaerobic growth (stationary phase). Initially, *ΔnahK* utilizes more nitrite than wild-type, but over time nitrite utilization stagnates in the mutant while steadily continuing in the wild-type. **(E)** NO concentration as a function of time. We observe initial accumulation of NO at early exponential phase (4 h) and then dissipation by stationary phase (16 h) in *ΔnahK*, while in contrast, NO is steadily produced in the wild-type strain. *p* values were calculated using unpaired, t-tailed t-test comparing *ΔnahK* to Wild-type, normalized for OD differences. \*\*\**p* value < 0.005, \*\*\*\**p* value < 0.001.

To measure the activity of NIR, we added nitrite to rich media under anoxic conditions, rather than nitrate, to bypass NAR. Under the conditions of this experiment, although *ΔnahK* has an early advantage, over the time course of the experiment, wild-type utilizes nitrite faster than *ΔnahK* (Figure 3C, 3D), indicating greater activity of NIR. Of note, the Δ*nahK* strain does not grow as well as wild-type under these conditions, thus nitrite utilization levels were normalized to OD.

We also measured NO accumulation as an additional measurement of NIR activity, as NIR converts nitrite to NO. The presence of NO was measured using an NO detection reagent called Daf-2 DA (24). As expected, in *ΔnahK*, there is an initial accumulation of NO in early exponential phase, however over time the amount of NO measured in the *ΔnahK* strain is significantly reduced (Figure 3E). By contrast, in the wild-type strain, NO accumulates throughout the course of the experiment. This reduction in NO produced by *ΔnahK* is consistent with a reduction in NIR activity and an overall lack of anaerobic respiration after early exponential phase.

All these data relating to reductase activity, taken together with our observed downregulation of reductase expression (Figure 2) in Δ*nahK*, leads us to conclude sustained denitrification is deficient in this strain. This suggests that NahK may be regulating denitrification under anaerobic conditions. We hypothesize this may be because NahK plays a role in reducing accumulation of toxic NO in *P. aeruginosa*.

### nahK is required for cell elongation under anaerobic conditions

At high concentrations, NO is toxic; bacteria that utilize denitrification have evolved methods to limit NO production and convert it to less toxic species (9-10). During anaerobic respiration, *P. aeruginosa* undergoes a dramatic morphological change and becomes highly elongated, compared to aerobically grown bacteria (26). This elongation has been attributed to diluting and dissipating accumulated NO. Further, elongation is required for robust biofilm formation under denitrifying conditions (26). Consistent with this, PAO1 *Δnir* mutants are unable to elongate or form biofilms under anaerobic conditions (24, 26).

To assess if changes in NO levels influence elongation in *ΔnahK*, we quantified cell elongation at several time points. Consistent with previous reports, we observed wild-type elongation under anaerobic conditions (Figure 4, 5). In contrast, *ΔnahK* initially elongates (after 4 h), but then reverts to normal cell morphology by stationary phase (16 h). This phenotype reverts when *ΔnahK* is transformed with a plasmid expressing *nahK*, but not an empty plasmid. These observations are consistent with the NO concentrations we observed in these strains (Figure 3).

**Figure 4.**
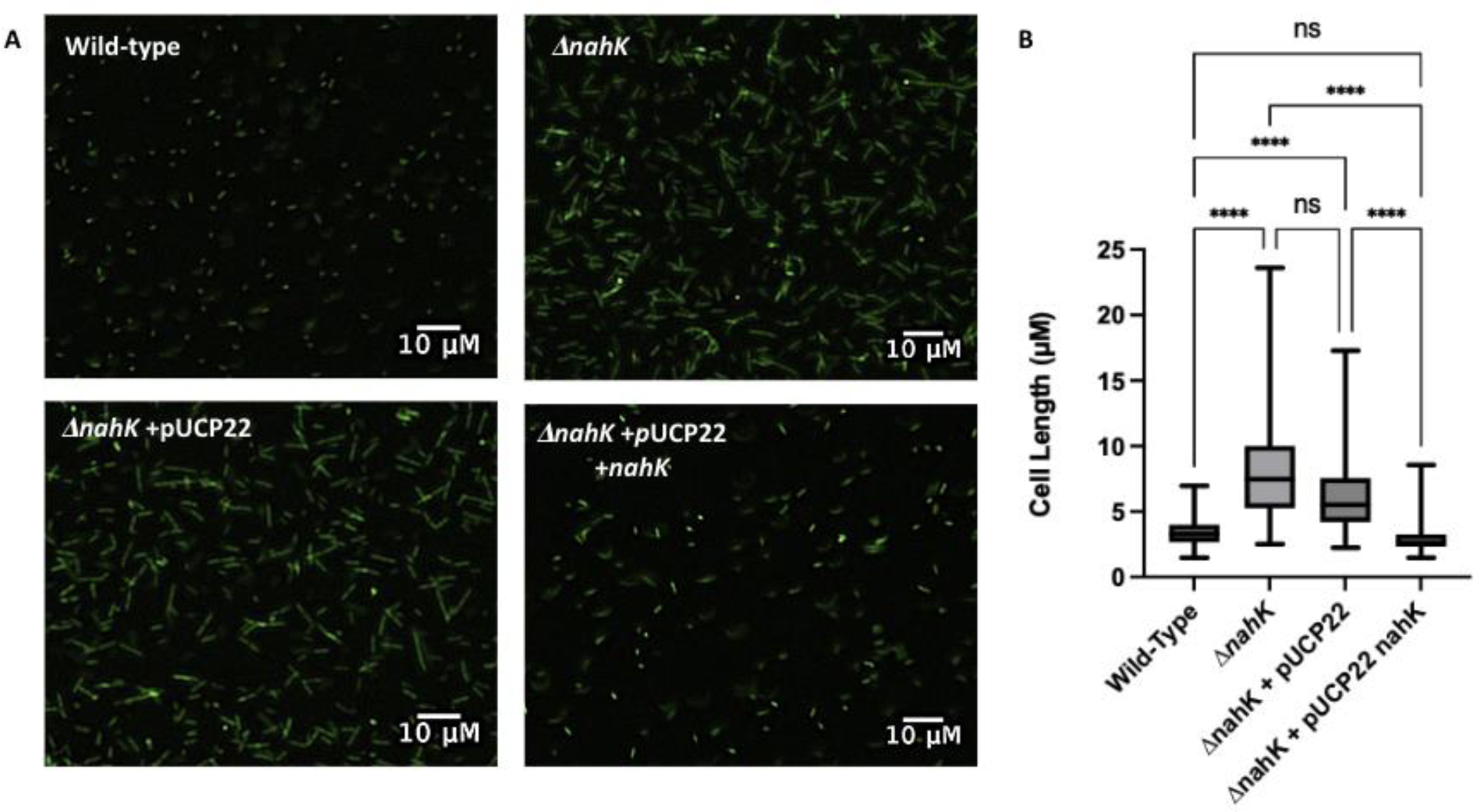
NO-dependent cell elongation is mis-regulated in *ΔnahK*. **(A)** Micrograph of Wild-type and *ΔnahK* in early exponential stage (4 h) of anaerobic growth. Bacteria were grown under anaerobic conditions supplemented with nitrate for 4 h. The premature elongation in *ΔnahK* is consistent with an initial increase in denitrification activity and NO accumulation. This phenotype reverts when *ΔnahK* is transformed with pUCP33 expressing *nahK*, but not empty pUCP33. Three independent experiments were carried out and representative images are shown. Cells were fixed in 4% paraformaldehyde and stained with Syto9. (**B)** Quantification of cell length. The select line tool was used to determine the cell length from scaled images using ImageJ. n=75; *p*-values were calculated using one-way analysis of variance and a Tukey multiple comparisons test. \**p* ≤ 0.5.

**Figure 5.**
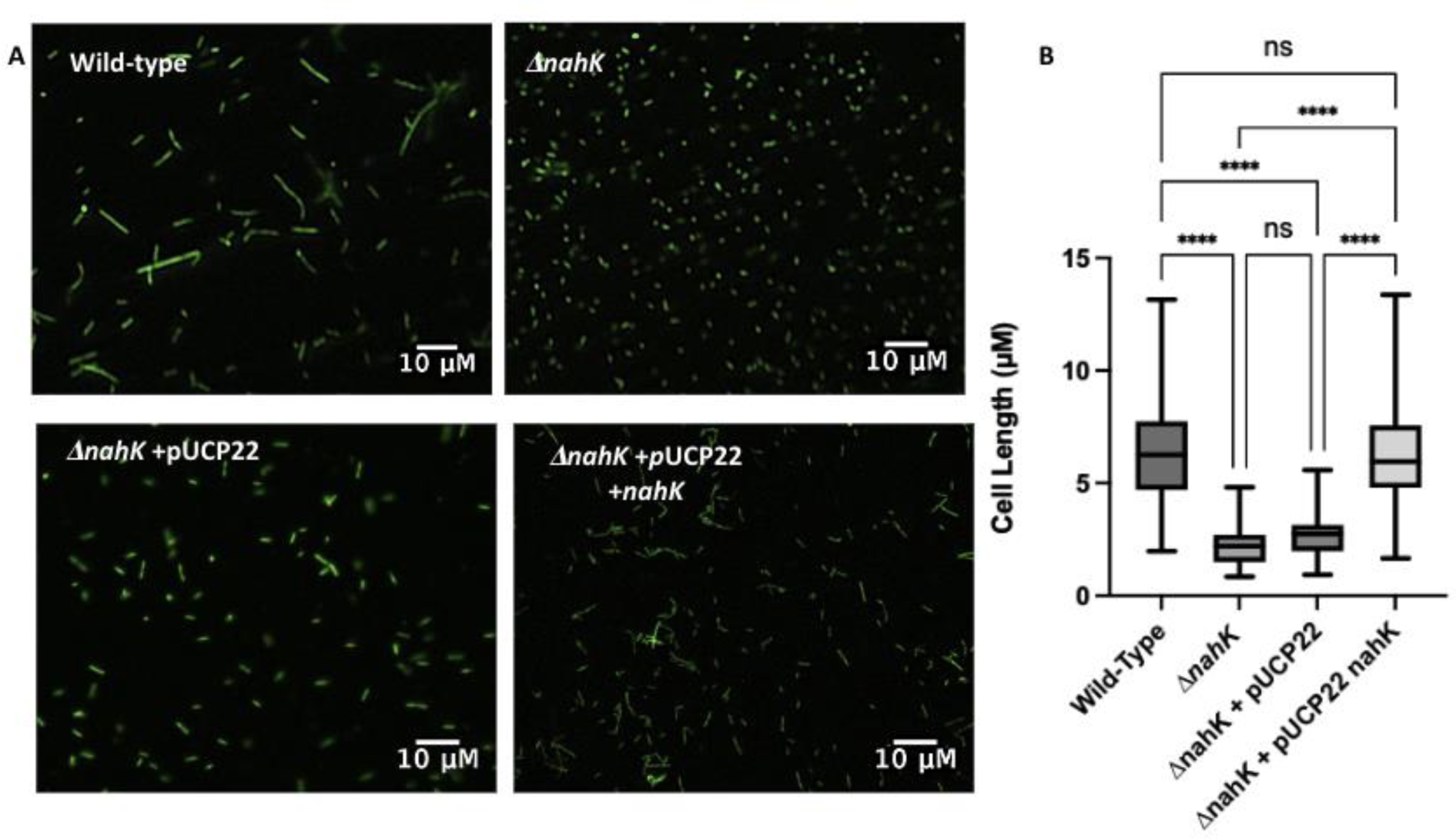
NO-dependent cell elongation is mis-regulated in *ΔnahK*. **(A)** Micrograph of Wild-type and *ΔnahK* during stationary phase (16 h) of anaerobic growth. Bacteria were grown under anaerobic conditions supplemented with nitrate for 16 h. Wild-type elongates under these conditions, however the early elongation observed in *ΔnahK* is absent at 16 h, suggesting that dentification activity and NO accumulation is reduced in this mutant. Elongation is partially recovered when *ΔnahK* is transformed with pUCP33 expressing *nahK*, but not empty pUCP33. Three independent experiments were carried out and representative images are shown. Cells were fixed in 4% paraformaldehyde and stained with Syto9. **(B)** Quantification of cell length. The select line tool was used to determine the cell length from scaled images using ImageJ. n=75; *p*-values were calculated using one-way analysis of variance and a Tukey multiple comparisons test. \**p* ≤ 0.5.

### Cell elongation is due to differences in NO accumulation

To confirm that the changes in elongation in *ΔnahK* are due to differences in NO accumulation over time, *ΔnahK* was grown in the presence of the NO scavenger carboxy-PTIO. After 4 h of growth under anaerobic conditions, *ΔnahK* no longer elongates in the presence of the NO scavenger (Figure 6). This supports our hypothesis that the early onset of elongation in the deletion strain is due to NO accumulation in early exponential phase (Figure 6).

**Figure 6.**
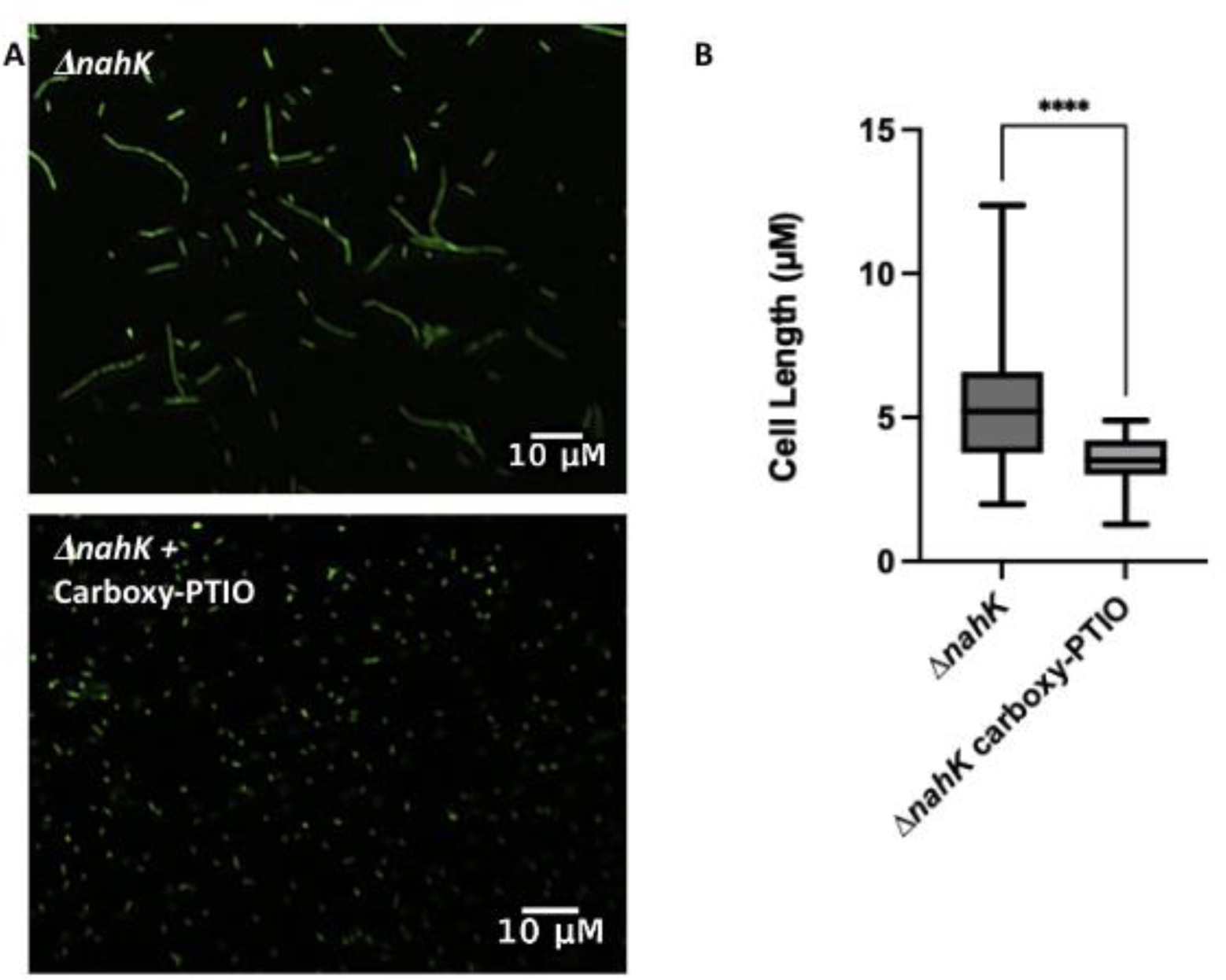
NO-dependent cell elongation is mis-regulated in *ΔnahK*. **(A)** Micrograph of *ΔnahK* grown in the presence and absence of NO scavenger Carboxy-PTIO. Bacteria were grown to early exponential phase (4 h) anaerobically with or without 2 mM NO scavenger Three independent experiments were carried out and representative images are shown. Loss of elongation in *ΔnahK* suggests that elongation during exponential phase is due to accumulation of NO. **(B)** Quantification of cell length. The select line tool was used to determine the cell length from scaled images using ImageJ. n=75; *p*-values were calculated using one-way analysis of variance and a Tukey multiple comparisons test. \**p* ≤ 0.5.

Likewise, to determine if the lack of cell elongation in *ΔnahK* during stationary phase is due to a reduced amount of NO produced later in growth, we grew *ΔnahK* in the presence of the long-acting NO donor DETA-NONOate until stationary phase. As expected, with the addition of exogenous NO, *ΔnahK* was elongated after 16 h of growth (Figure 7). In wild-type *P. aeruginosa*, although NOR is reducing NO to N_2_O, there is an overall accumulation of NO under anaerobic conditions contributing to the sustained cell elongation over time (26). These results suggest that the lack of accumulation of NO during stationary phase is contributing to this loss of elongation in *ΔnahK*.

**Figure 7.**
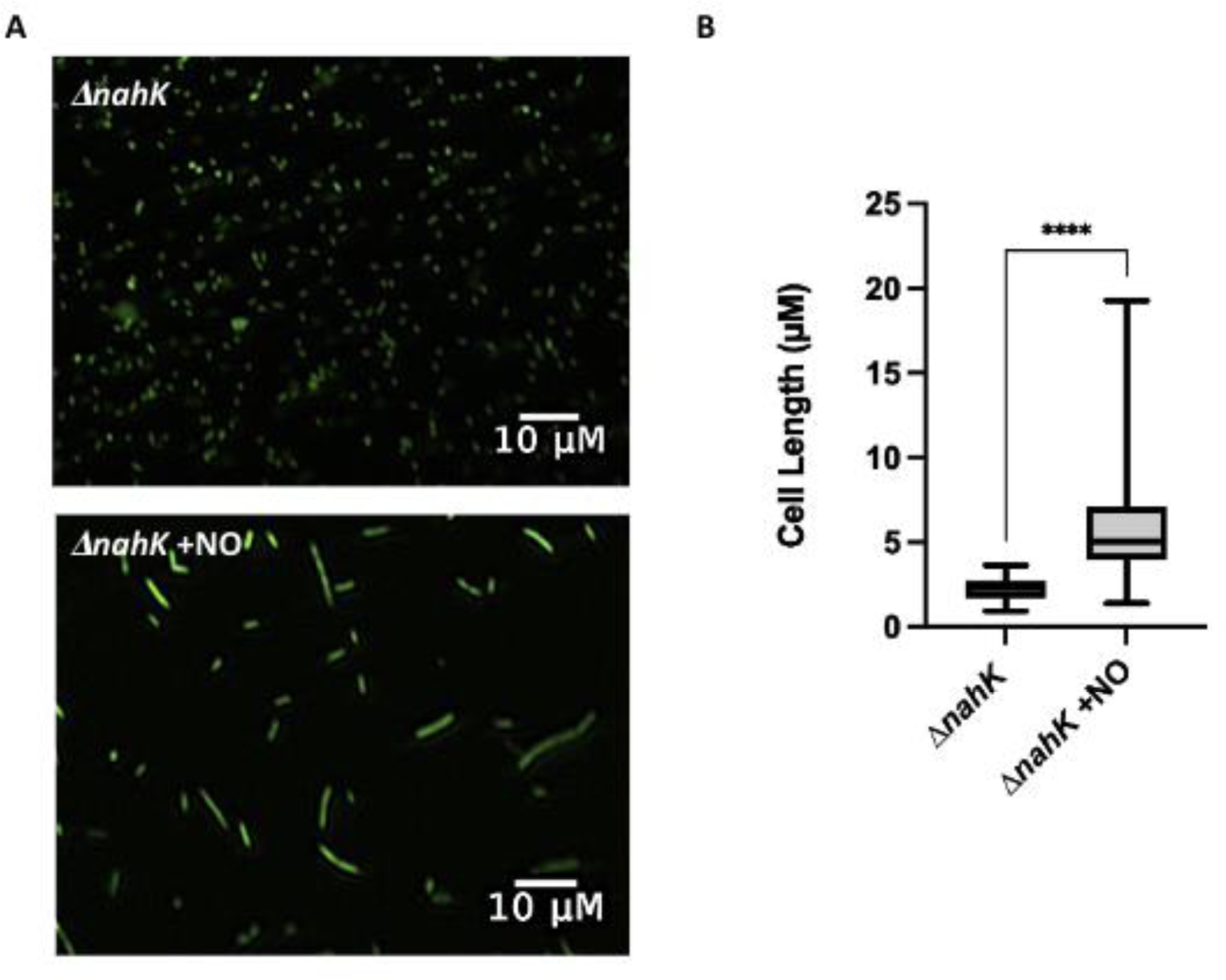
NO-dependent cell elongation is mis-regulated in *ΔnahK*. **(A)** Micrograph of *ΔnahK* grown in the presence and absence of exogenous NO. Bacteria were grown to early exponential phase (4 h) anaerobically with or without 100 µM DETA-NONOate Three independent experiments were carried out and representative images are shown. Recovery of elongation in stationary phase in ΔnahK in the presence of NO supports a reduction in NO accumulation at this timepoint **(B)** Quantification of cell length. The select line tool was used to determine the cell length from scaled images using ImageJ. n=75; *p*-values were calculated using one-way analysis of variance and a Tukey multiple comparisons test. \**p* ≤ 0.5.

### Loss of NO accumulation and elongation in ΔnahK may be due to reduced RsmA activity

NahK is one of several kinases in the GacS multikinase network that are important in regulating RsmA (16). RsmA is a posttranscriptional regulator that binds mRNA sequences encoding proteins involved in biofilm formation, thereby preventing their translation and promoting bacteria motility (27). In our previous work, we have shown that overexpression of *rsmA* in *ΔnahK* restored pyocyanin levels back to wild-type levels, suggesting the main effect of deletion of *nahK* is the reduction of RsmA levels (21). To determine if the denitrification phenotypes we observed in this study are due to a similar mechanism, we measured cell elongation *ΔnahK* expressing *rsmA* during stationary phase under anaerobic conditions (Figure 8). As expected, the overexpression of *rsmA* restored elongation in *ΔnahK,* supporting our previous conclusion that RsmA levels are inhibited in *ΔnahK.* The data presented here suggest a novel role for NahK and the RsmA network in the regulation of denitrification activity and accumulation of NO levels.

**Figure 8.**
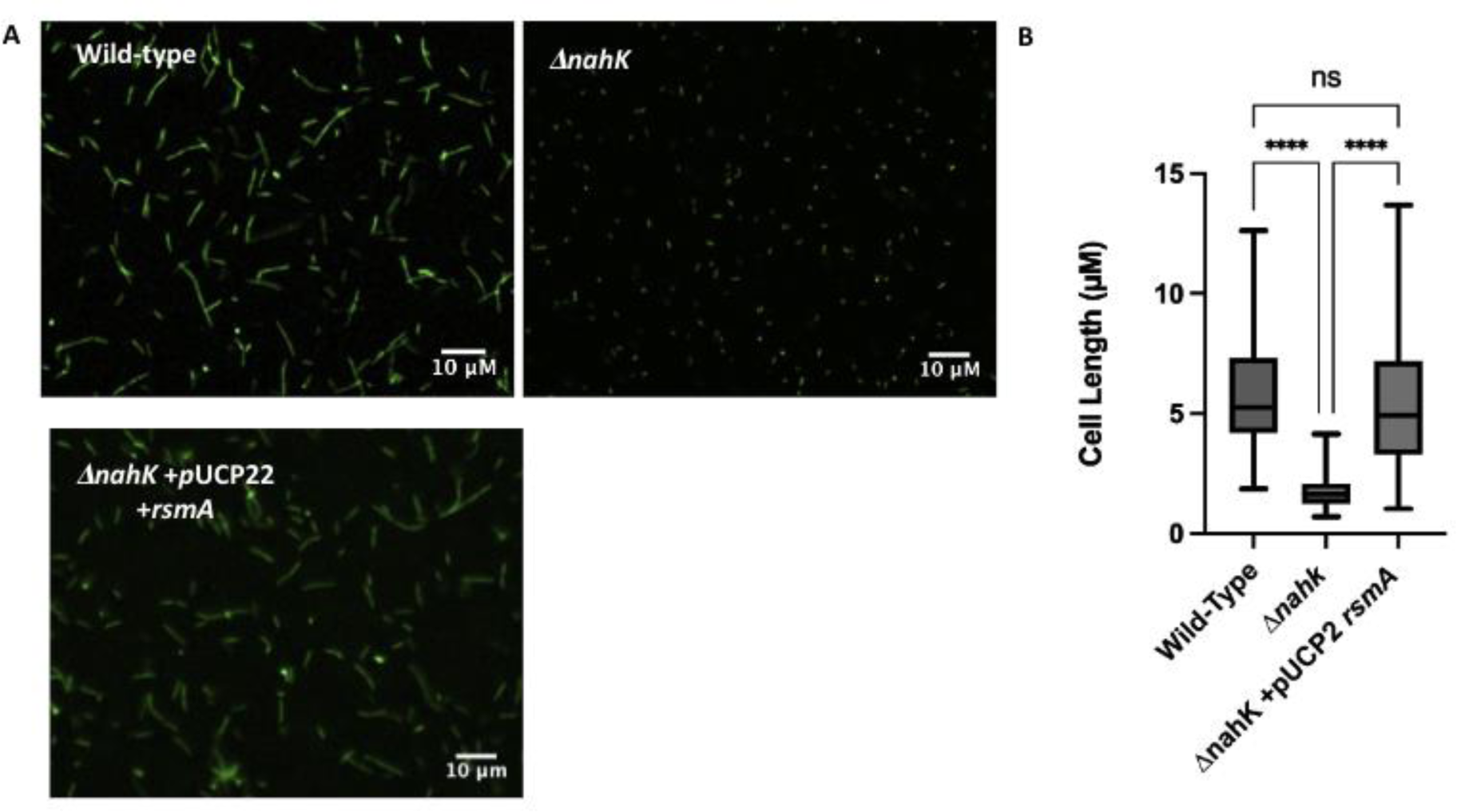
NahK regulates NO-dependent cell elongation through modulation of RsmA activity. **(A)** Micrograph of overexpression of RsmA in *Δnahk.* Bacteria were grown under anaerobic conditions supplemented with nitrate until stationary phase (16 h). Three independent experiments were carried out and representative images are shown. Overexpression of *rsmA* in Δ*nahK* restores elongation to levels similar to that seen in the wild-type. These data suggest that RsmA levels are reduced in *ΔnahK*, and that the reduction of NO accumulation in Δ*nahK* is in part due this reduction in RsmA activity. **( B)** Quantification of cell length. The select line n=75; *p*-values were calculated using one-way analysis of variance and a Tukey multiple comparisons test. \**p* ≤ 0.5.

### NahK promotes biofilm formation under anaerobic conditions

Anaerobic cell elongation, presumably due to accumulated NO, has been linked to better anaerobic biofilm formation in *P. aeruginosa* (25). Because in *ΔnahK* we observe a lack of cell elongation during stationary phase, and a reduction of NO accumulation, we decided to investigate anaerobic biofilm formation in this mutant. For a baseline comparison, we first compared *ΔnahK* and wild-type in an aerobic static biofilm assay (Figure 9). We observed an increase in biofilm formation in *ΔnahK*, compared to wild-type, under these conditions. Under anaerobic conditions, however, we observed a significant increase in wild-type biofilm formation, when compared to biofilm formation under aerobic conditions. This change in biofilm formation as a function of oxygen concentrations is not observed in *ΔnahK*. Furthermore, overexpression of *rsmA* in Δ*nahK* restores the mutant to wild-type biofilm formation under both aerobic and anaerobic conditions. We attribute the reduction in anaerobic biofilm formation in *ΔnahK* to the loss of cell elongation and NO accumulation in this mutant, downstream of reduced RsmA levels.

**Figure 9.**
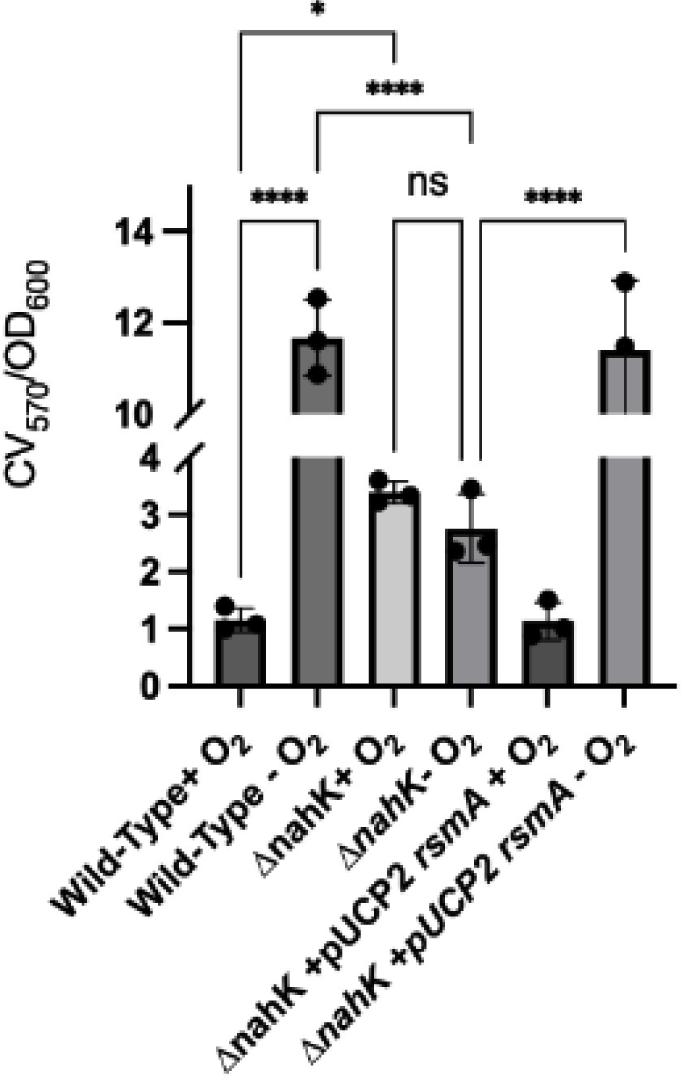
Anaerobic biofilm formation is mis-regulated in *ΔnahK*. A static biofilm assay was utilized to assess biofilm formation. Anoxic incubation was achieved using Thermo Scientific™ AnaeroPack™-Anaero Anaerobic Gas Generator system. Surface attached biofilms were stained using 0.1% crystal violet (CV) solubilized in 30% acetic acid. Biofilm mass was quantified by the absorption of CV (570 nm) normalized to cell optical density (600 nm). *p*-values were calculated using one-way analysis of variance and a Tukey multiple comparisons test. \**p* ≤ 0.5.

Presumably NahK activity is modulated by the NosP/NO complex in the wild-type, which directs RsmA-mediated biofilm regulation in response to NO generated during denitrification. Interestingly, under aerobic conditions, NO detection triggers biofilm dispersal (15), however, NO-driven cell elongation is required for robust biofilm formation when oxygen is not present (26), thus it may be advantageous for bacteria to prevent NO-mediated biofilm dispersal under anaerobic conditions. Likewise, RsmA activity typically promotes bacteria motility, however, these data suggest that under anaerobic conditions, RsmA promotes biofilm formation due to changes in response to NO accumulation. NahK may contribute to this activity switch. Understanding precisely how and why NO/NosP/NahK contributes to RsmA and quorum sensing mediated biofilm regulation in the presence and absence of oxygen is needed to better understand the role of NO in biofilm formation. More experiments will be required to understand this fully.

## Discussion

Here we describe the role of the histidine kinase NahK on NO-dependent regulation of denitrification under anaerobic conditions (Figure 10). We have shown that loss of *nahK* results in early and unsustainable transcription-driven denitrification activity, with accompanying cell death and loss of NO-driven cell elongation at higher cell densities. Presumably the early generation of NO is responsible for the subsequent downregulation of the denitrification machinery, possibly as part of a feedback mechanism to prevent toxic levels of NO from accumulating. Loss of NO-driven cell elongation also results in a reduction of biofilm under oxygen-limited concentrations. RsmA overexpression in the Δ*nahK* mutant restores wild-type cell elongation and biofilm phenotypes. These data therefore suggest that NahK activity regulates RsmA levels and that RsmA is required for regulation of denitrification.

**Figure 10.**
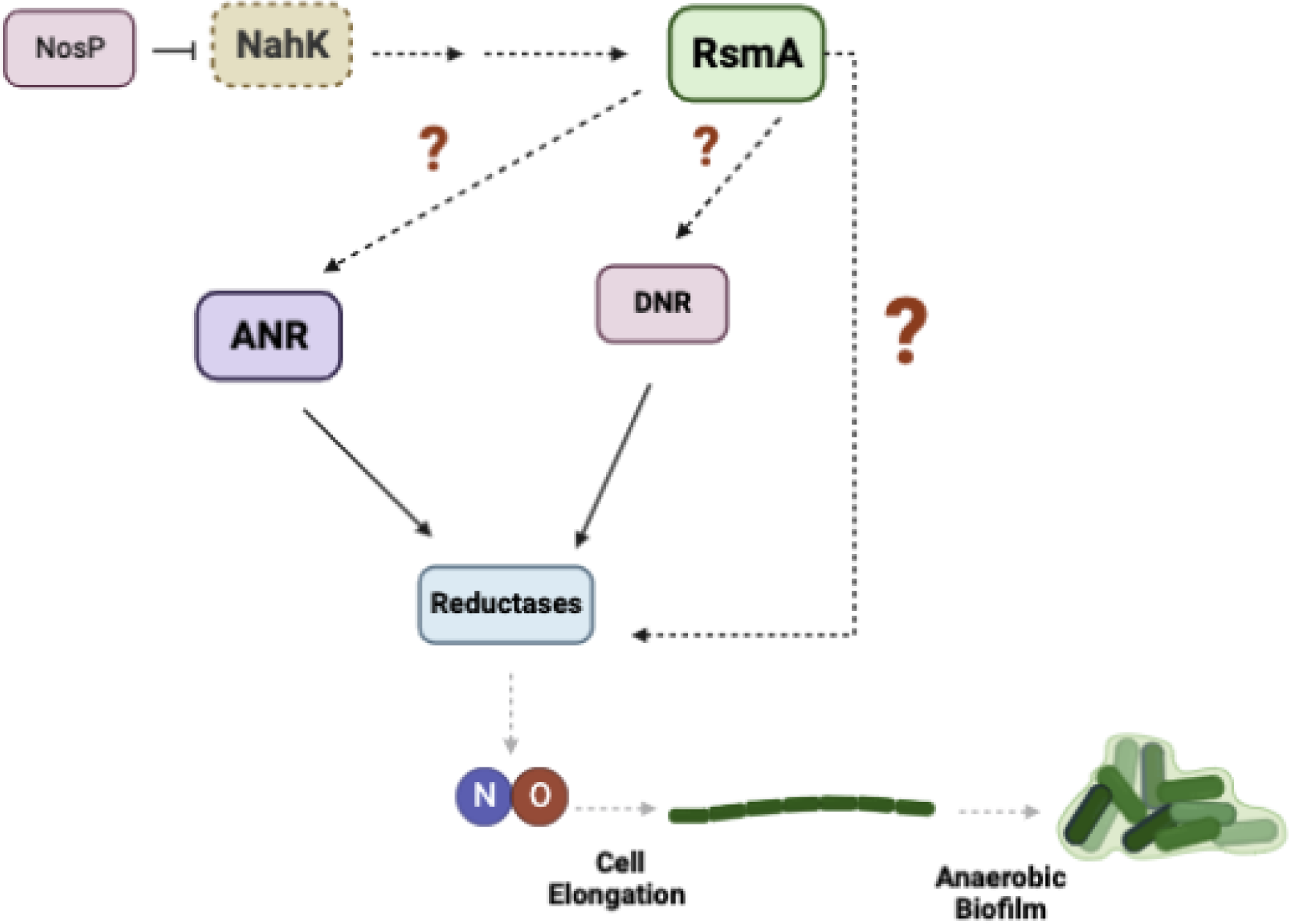
NahK regulates denitrification and anaerobic biofilm formation through the modulation of RsmA levels. Inhibition of NahK contributes to a reduction in RsmA signaling. This reduction reduces the expression of the reductases and reduces denitrification activity. This loss of denitrification activity contributes to reduced levels of NO accumulation resulting in a loss of cell elongation and biofilm formation under anaerobic conditions.

Given that NosP binds NO and regulates NahK (15), it would make sense that NO generated during denitrification is acting as a feedback signal to regulate RsmA through NahK. However, it is clear that NahK activity is not needed for detection of the NO levels that drive cell elongation. The *ΔnahK* strain elongates in response to NO generated at low cell density (Figure 4) and upon supplementation with exogenous NO at high cell density (Figure 7). We hypothesize that loss of NahK might cause the cells to be more sensitive to low concentrations of NO than wild-type. The amount of NO added to *ΔnahK* that restores cell elongation at high cell density under anaerobic conditions is actually very low (∼100 nM), an amount that is insufficient to drive elongation or other stress phenotypes when added to the wild-type strain (Supplemental Figure 2). Interestingly, when *ΔnahK* is grown aerobically in the presence of NO, we do observe some elongation, which is not seen in the wild-type under these conditions (Supplemental Figure 2), but not to the same extent as under denitrifying conditions; this may also be evidence that *ΔnahK* is more sensitive to NO than wild-type. Exactly why the cells are more sensitive to NO is not clear at this point.

Nonetheless, there are data in the literature that may help to explain the results we report here. *Burkholderia pseudomallei* can generate intracellular NO through denitrification under low oxygen conditions (28). The presence of nitrate and nitrite has been shown to reduce pellicle biofilm formation in *B. pseudomallei,* both aerobically and anaerobically (29). Further, in the presence of nitrate and nitrite, the *nosP/nahK/narR* operon is upregulated in *B. pseudomallei*, along with several other genes involved in anaerobic metabolism and antibiotic tolerance (28). This upregulation is dependent on the two-component system NarX/NarL which is also a conserved two-component system in *P. aeruginosa* (30-31). NarX is sensitive to nitrate and initiates the activation of NarL through autophosphorylation (30-31). This suggests that NarXL may regulate NosP and NahK in *B. pseudomallei*, and perhaps also in *P. aeruginosa*. Taken together, these studies suggests that detection of nitrate is essential for regulation of metabolism and biofilm formation, possibly through NahK/RsmA-regulated fine tuning of intracellular NO levels. Further studies will be needed to test this mechanism and determine if it is conserved in other facultative aerobic bacteria.

It is surprising that the role of the RsmA network in denitrification remains relatively unexplored in *P. aeruginosa*. As mentioned above, RsmA is regulated by the GacS multikinase network, and is thought to serve as molecular switch between biofilm formation and bacterial motility (20). The GacS network includes GacS, RetS, SagS, PA1611, and NahK, which indirectly regulate RsmA activity through its inhibitory small regulatory RNAs *rsmY* and *rsmZ* (19). In *ΔrsmA*, a decrease in the expression of *anr*, *nirS*, *narG* and *norC* has been reported (27). Loss of *rsmY* has been previously reported to upregulate azurin, a blue copper protein that supports denitrification by transferring electrons to NIR and is expressed under low oxygen conditions (19, 32). Further, two studies have recently reported that nitrate reductase activity is reduced in *ΔretS* under anaerobic conditions (33-35). RetS is known to inhibit the type IV secretion system gene cluster H2-T6SS. One of the H2-T6SS secreted proteins is ModA, a molybdate scavenger, which is required for anaerobiosis because molybdenum is an important cofactor for NAR (33-35).

Moreover, RsmA is an inhibitor of *lasR*, which is an inhibitor of denitrification (11, 20). Interestingly, a common mutation in *P. aeruginosa* isolates from CF patients is a deletion of the *lasR* gene which is thought to promote a survival in anaerobic biofilms in the CF lung (36). This increase in fitness is due to a shift in metabolism in these mutants in which there is less consumption of oxygen and more utilization of nitrate (36). This shift in metabolism is attributed in part to increased activity of ANR and the denitrification reductases, although the molecular mechanism of the increase in ANR activity is currently unknown (37). This metabolism shift in the *ΔlasR* mutant also promotes fitness by promoting higher antibiotic resistance to tobramycin and ciprofloxacin, which are commonly used to treat *P. aeruginosa* infections in CF patients compared to its LasR+ counterpart (37,38). Interestingly, these studies lead to the prediction that NahK may have implications in antibiotic resistance. All these observations are important in electron shuttling under microaerobic conditions and the changes in their production allude to a greater role of the GacS multikinase network in denitrification regulation although, each of these factors need to be individually characterized to establish the overall mechanism of this system.

Furthermore, we speculate that there may be a connection between modulation of denitrification by NahK and the formation of persister cells under anaerobic conditions (e.g., in a biofilm). Persister cells are a subpopulation of metabolically inactive bacteria within a microbial community that are tolerant to harsh conditions, including antibiotics, and help ensure the long-term survival of a biofilm. Biofilms promote the formation of persister cells in part due to the heterogeneity of oxygen and nutrient availability within the biofilm (39). In this study we demonstrated that cells lacking NahK have mis-regulated denitrification and appear to overreact to NO produced during denitrification, eventually leading to stagnated respiration. In the absence of oxygen and with minimal denitrification activity, it is not clear how the *ΔnahK* cells survive without going into a dormant or persistent state. To be clear, we do observe some cell death (Figure 1), but it is relatively minimal, so it seems that most of the cells survive with very minimal respiration in our experiments.

We are not sure how persistence may occur in our experiments, but NO can activate the SOS response upon DNA damage (8), which is one of many factors contributing to the formation of persister cells. RecA de-represses genes involved in DNA repair (40). The activation of RecA also contributes to the formation of membrane vesicles, which are involved in extracellular transport of proteins and quorum sensing signals, in addition to playing a role in antibiotic neutralization (41-42). It has been reported that *P. aeruginosa ΔnirS* is unable to produce membrane vesicles anaerobically, suggesting that production of NO is essential for the activation of SOS and formation of these vesicles (8). If, as we have speculated, the *ΔnahK* strain is sensitive to NO, or just more stressed, the amount of NO produced by denitrification at low cell density may be triggering an SOS response.

An additional possible link between NahK and persister cell formation is based on the role of NahK in regulating quorum sensing and PQS (16, 21). Exogenous addition of the PQS precursor 2-AA promotes the formation of persister cells. When *mvfR*, the gene that encodes the protein that produces 2-AA, is deleted from *P. aeruginosa*, fewer persister cells are formed than in the wild-type strain. Addition of 2-AA to *ΔmvfR* results in an increase in a ribosomal modulation factor which promotes ribosomal inactivity in persister cells (43). We have previously shown that PQS is overproduced in *ΔnahK* (43, 21). This suggests there may be a connection between NahK activity and regulation of levels of 2-AA contributing to persistence, virulence, and antibiotic tolerance.

In conclusion, our study implicates NO and NahK in regulating denitrification through RsmA. Understanding how NahK directs dentification activity may have important implications for how NO sensing through NosP contributes to the molecular mechanisms that regulation biofilm formation and possibly persistence.

## Materials and Methods

### Bacterial strains and growth conditions

Bacterial strains used in this study are described in Supplemental Table 1. *P. aeruginosa* strains were grown aerobically in Luria-Bertani (LB) Broth. Anaerobic bacterial growth was achieved using Thermo Scientific™ AnaeroPack™-Anaero Anaerobic Gas Generator system or in a Coy vinyl anaerobic chamber. Planktonic bacteria were grown in Hungate anaerobic tubes. The gas composition inside the anaerobic chamber used a mix of nitrogen and hydrogen (95% nitrogen and 5% hydrogen, respectively). To support anaerobic growth, 25 mM NaNO_3_ or 25 mM NaNO_2_ was added to the media. To confirm growth conditions were anaerobic, no detectable growth was observed in LB without NaNO_3_. Cultures were grown at 37^°^C shaking at 250 rpm.

### Growth Curve and CFU counting

Day cultures were grown from overnight cultures to an OD_600_ of 1.0 in LB media. Cultures were diluted to an OD_600_ of 0.005 into 5 mL of fresh LB or into 5 mL of fresh LB supplemented with 25mM NaNO_3_. Cultures were grown both aerobically and anaerobically. Every two hours, the OD_600_ was measured and cultures were returned to the incubator. CFU counting was conducted using a previously described method (44).

### Quantitative PCR (qPCR)

For qPCR, anaerobic conditions were grown in 5mL LB media supplemented with 25 mM NaNO_3_ for four hours (exponential phase) or sixteen hours (stationary phase). For aerobic conditions, bacteria were grown in 5mL LB media to an OD_600_ of 1.0. For all conditions, bacteria were grown at 37°C with agitation. Total cellular RNA extraction was performed using the RNeasy Mini Kit (Qiagen). RNA yield was quantified using nanodrop and purity was confirmed by using a 1% agarose gel. cDNA synthesis was performed using RevertAid First Strand cDNA synthesis kit (Thermo Scientific) from purified RNA. qPCR reaction included 0.3µM of forward and reverse primer described in **Supp. Table 2** equal amounts of synthesized cDNA and 5µL OF SYBR green master mix (Thermo Scientific) for a total reaction volume of 10 µL. Reaction was performed on a LightCycler 480 with the cycler parameters of 95°C for 10 min, then 40 cycles of 95°C for 15 s and 60°C for 60s. GyrA was used as a housekeeping gene and relative expression was determined using a standard curve (45).

### Denitrification Activity Assays

Denitrification activities were measured using previously described methods (24) NO_2_ concentration was determined using a Greiss Detection Kit (Thermo Scientific). To assess NO_2_ concentration, bacteria were grown anaerobically in 5mL LB media supplemented with 25 mM NaNO_3_ or 25mM NaNO_2_ to measure NO_2_ production and NO_2_ utilization over time respectively. Supernatant was collected from centrifuged cells and incubated with prepared Griess reagent for 30 minutes. Absorbance was read at 548 nm in reference to the photometric reference sample. Concentration was assessed using a standard calibration curve of NaNO_2_.

### NO Detection

Cellular NO levels were measured using a previously described method (24) using the NO detection reagent diaminofluorescein-2 diacetate (DAF-2 DA).1mL of shaking cultures grown under anaerobic conditions were incubated with 10 μM DAF-2 DA at 7°C for 1 h and washed with PBS. Fluorescence of the reaction product, DAF-2 T was assessed using a SpectraMax iD3 plate reader at an excitation wavelength of 495 nm and emission wavelength of 515 nm normalized to OD_600_.

### Fluorescent microscopy and cell length quantification

For all sample preparations, cells were fixed in 4% paraformaldehyde. The nucleic acid stain SYTO9 green fluorescent dye was used at a final concentration of 10 µM. A Zeiss Axio Vert.A1 inverted epifluorescence microscope equipped with a Lumencor Sola Light Engine, a Lumenera 8MP Infinity3 Camera, a Zeiss GFP fluorescence filter set to 38 HE, and a Zeiss A-Plan 40× N.A. 0.55 objective was used. The select line tool was used to determine the cell length from scaled images using ImageJ. To assess cell elongation in the presence of exogenous NO, 100µM of DETA-NONOate was added to LB media supplemented with 25 mM NaNO_3_ where cultures were grown anaerobically for 16 hours. To assess cell elongation dependence on the presence of NO, 2mM of the NO scavenger carboxy-PTIO was incubated with *ΔnahK* for four hours under anaerobic conditions.

### Static Biofilm Assay

Ability of wild-type and *ΔnahK* to form biofilm under aerobic and anaerobic conditions was assessed using a modified microtiter dish assay in 6-well plates to quantify surfaced attached bacteria grown statically (46). To assess biofilm formation aerobically, 5mL day cultures were prepared from overnight cultures and grown until the cultures reached an OD_600_ of 1.0. 500µL of day culture were added to each well which contained 9.5mL of LB or LB supplemented with 25mM NaNO_3_. To assess biofilm formation anaerobically, day cultures at an OD_600_ 1.0 were diluted to an OD_600_ of 0.005 into 5mL of fresh LB and grown for 16 h. 500 µL of anaerobic overnight culture was added to each well which contained 9.5mL of LB supplemented with 25mM NaNO_3_ to achieve anaerobic growth, Thermo Scientific™ AnaeroPack™-Anaero Anaerobic Gas Generator system was used.6-well plates were incubated at 37 °C statically for 16 h. After 16h, OD_600_ of the planktonic cells was recorded on the SpectraMax iD3 plate reader. The remaining planktonic cells were removed and the plate was rinsed with ultrapure water. The plate was dried and stained with 2mL/well 0.1% crystal violet (CV) for 15 minutes with agitation. Excess CV was rinsed with ultra-pure water. After drying for 1h, 2mL 30% acetic acid was added to each well to solubilize the CV. CV was quantified by absorbance and was read at 570 nm to quantity CV. CV570 was normalized to OD_600_ to account for variation in planktonic growth although OD_600_ was consistent from strain to strain in each condition.

## Data Availability

All requests for resources and reagents should be directed to and will be fulfilled by the Lead Contact Elizabeth Boon. All constructs and cell lines generated in this study are freely available upon request.

---

**Supplemental Figure 1.**
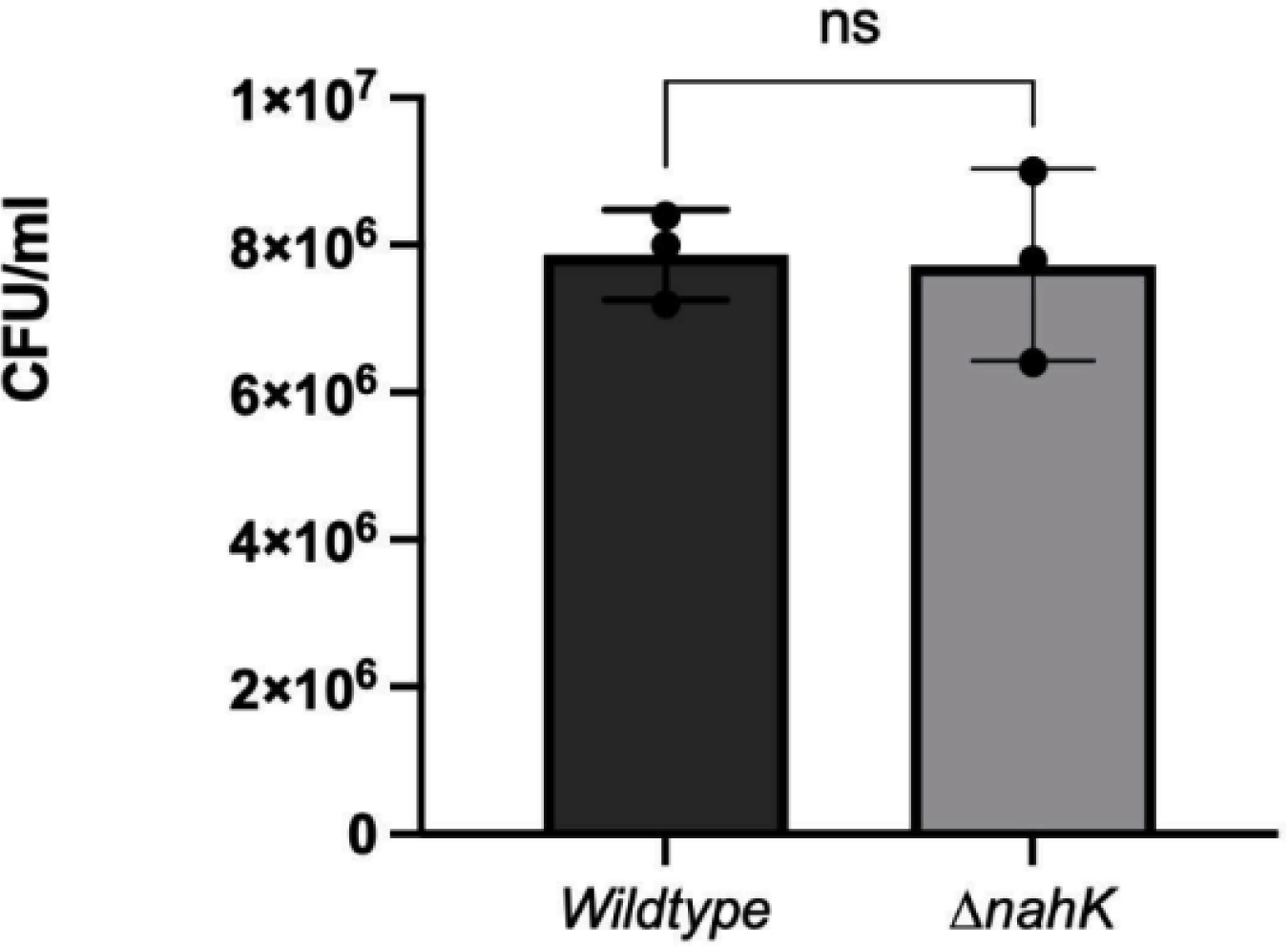
CFU Counting at OD 1.0 under aerobic shaking conditions. The values represent means ± standard deviations from 3 independent experiments. p values were calculated using unpaired, t-tailed t-test comparing Δ*nahK* OD to Wild-type OD. \**p* ≤ 0.5.

**Supplemental Figure 2.**
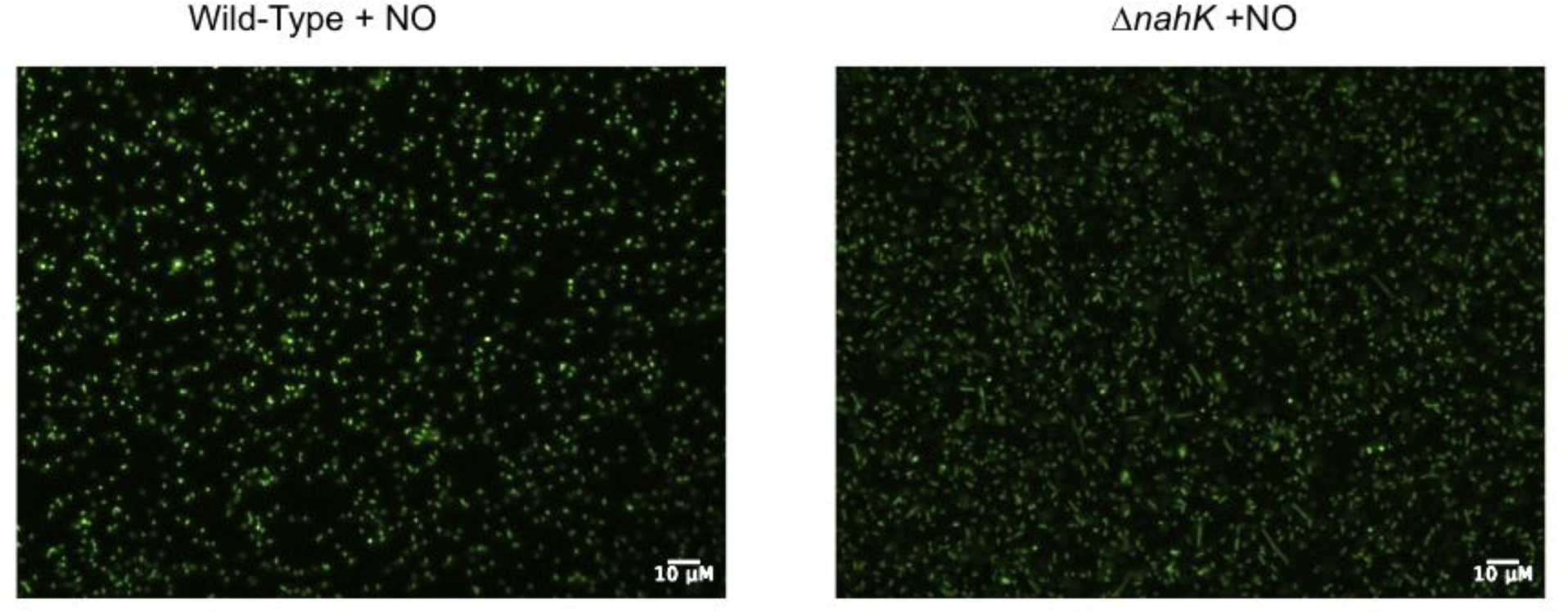
Micrograph of wild-type and *ΔnahK* grown in the presence NO aerobically. Bacteria were grown for sixteen hours aerobically with 100 µM DETA-NONOate. Three independent experiments were carried out and representative images are shown.

**Supplemental Table 1.** Strains used in this study.

| Strains | Relevant Characteristics | Source |
| --- | --- | --- |
| ***P. aeruginosa*** |  |  |
| UCBPP- PA14 | Wildtype | Lab Collection |
| PA14 $\Delta nahK$ | PA14 deletion mutant for PA14_38970 (*nahK*). | Lab Collection (20) |

**Supplemental Table 2.** Plasmids used in this study.

| Plasmid | Relevant Characteristic | Source |
| --- | --- | --- |
| pUCP22 | Amp<sup>R</sup> Gen<sup>R</sup> | Addgene |
| pUCP22 *nahK* | Amp<sup>R</sup> Gen<sup>R</sup> containing *nahK* gene. | Lab Collection (20) |
| pUCP22 *nahK* | Amp<sup>R</sup> Gen<sup>R</sup> containing *rsmA* gene. | Lab Collection (20) |

**Supplemental Table 3.**
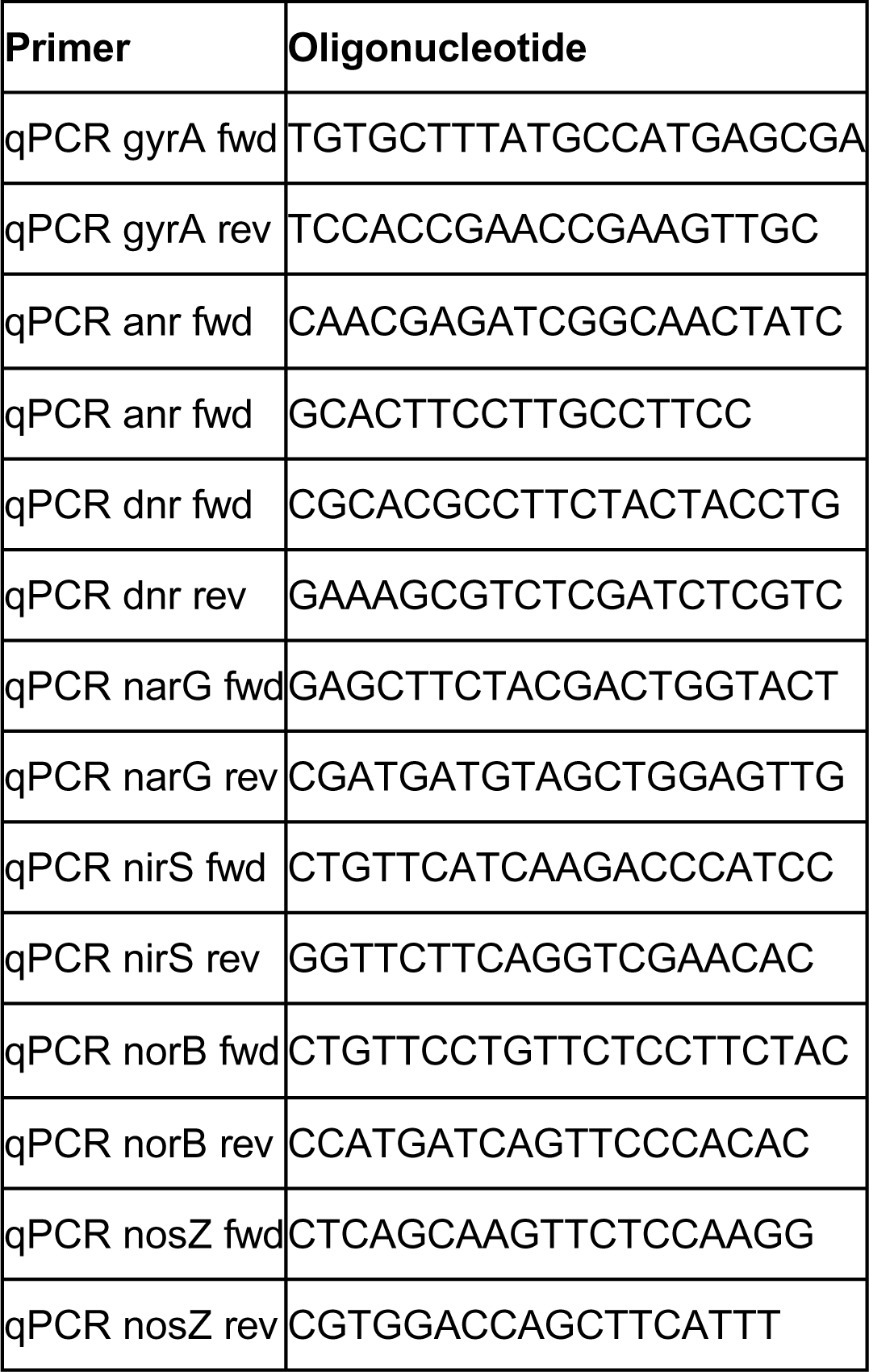
Primers used in this study.

